# Exploring simultaneous production of polyhydroxybutyrate and exopolysaccharides in cyanobacteria-rich microbiomes

**DOI:** 10.1101/2024.12.15.627710

**Authors:** Beatriz Altamira-Algarra, Joan García, Cristiana A. V. Torres, Maria A.M. Reis, Eva Gonzalez-Flo

## Abstract

The aim of this study was to explore the viability of the dual production of polyhydroxybutyrate (PHB) and exopolysaccharides (EPS) by seven microbiomes rich in cyanobacteria. Our initial experiments involved to screen for EPS-producing candidates and examine the impact of salinity and acetate on EPS synthesis. Salinity’s known influence on EPS production and acetate’s role in enhancing PHB production guided our parameter selection. Surprisingly, neither the introduction of an external carbon source (acetate) nor exposure to an abiotic stressor (salt) significantly altered EPS synthesis rates, which ranged from 25 to 150 mg·L^-1^, or its composition, with glucose being the dominant sugar component. Scaling up to a 3 L photobioreactor, we achieved simultaneous biopolymer production, reaching up to 205 mg·L^-1^ EPS and 87 mg·L^-1^ PHB. Additionally, the presence of uronic acid in the EPS facilitated biomass flocculation, streamlining the separation process, and potentially reducing associated time and costs.

## Introduction

As global attention turns towards sustainable biotechnological solutions, there is a rising interest surrounding the diverse range of high-value bioproducts derived from the cultivation of cyanobacteria, including pigments, polymers, and the biomass itself. Particularly, exopolysaccharides (EPS) and polyhydroxybutyrate (PHB) have attracted significant attention due to their biological and physico-chemical characteristics [1–4]. These properties position EPS as promising candidates for various applications as thickeners, stabilizers, and gelling agents within the agri-food, pharmaceutical, and cosmetics industries [5,6], while PHB can become a suitable replacement for petroleum-based polymers having potential applications in agriculture, food packaging or medicine [3,4,7]. However, the full potential of these bioproducts remains limited by economic issues which in turn are linked to gaps in fundamental knowledge of the processes [8].

PHB is accumulated intracellularly by cyanobacteria under nutrient-limited conditions. Trials with cyanobacteria wild-type (wt) strains monocultures in autotrophic conditions typically have achieved low yields, usually below 15% dry cell weight (dcw) PHB [9–12]. Various efforts have been attempted to develop new strategies to increase these yields and corresponding production, including molecular biology techniques, such as the introduction of genes associated with PHB metabolism to cyanobacteria cells [13–15], or supplementing cultures with an external organic carbon source. The addition of acetate (Ac) to the culture medium has resulted in PHB production of up to 46% dcw PHB in monocultures of *Anabaena* sp. [16], 26 %dcw PHB in *Synechoccocus* sp. [10] and 22 %dcw PHB in *Synechocystis* sp. [17].

Cyanobacteria produce EPS as part of their metabolism, playing crucial roles in cell adhesion, self-protection, and providing energy supply, particularly in extreme environments [18,19]. Comprising a variety of heteropolysaccharides, consisting of up to twelve different monosaccharides, these polymers form a protective barrier around the cell surface. In addition, they also incorporate diverse non-sugar substituents on their structure, including proteins, acyl or sulphate groups, or uronic acids, which contribute to their functional properties [19,20]. Cyanobacterial EPS can be categorized into two main groups: (i) those associated with the cell surface (known as cell-bound polysaccharides, CPS), (ii) and the polysaccharides released into the surrounding environment (referred to as released polysaccharides, RPS) [21]. Several cyanobacteria species, including *Synechococcus* sp., *Nostoc* sp., *Cyanothece* sp. or *Spirulina* sp. have been recognized as potential EPS producers [5,22]. However, comprehensive understanding of their synthesis remains challenging due to the strain-dependent responses to conditions in the culture affecting EPS synthesis. For example, abiotic stresses, such as light intensity, temperature, salinity, or pH, have shown diverse impacts on EPS production across various strains. Salinity, specifically, displays contrasting influences on EPS production among cyanobacteria species. It had either negative or no discernible impacts on EPS production in *Cyanothece* sp. [23,24], *Anabaena* sp. [25], *Spirulina* sp. [26] or *Aphanocapsa* sp. [27]. On the contrary, in other works, salinity stress positively influenced EPS production in *Synechocystis* sp. [28], *Synechococcus* sp. [29], *Oscillatoria* sp. [30] and *Cyanothece* sp. [31]. Similarly, the availability of macro and micronutrients, such as nitrogen, phosphorus, sulphate, or calcium, have also been explored to enhance EPS production, yielding strain-dependent outcomes and contradictory results [5,22].

The individual production of EPS and PHB by cyanobacteria monocultures has been studied [3,5,22,32]. Nevertheless, investigations into biopolymers synthesis within cyanobacterial microbiomes—a diverse microbial culture comprising various cyanobacterial strains and other microorganisms— have been limited to very few studies [33–35]. Notably, these studies have shown promising results, achieving for the first time in the field, PHB yields of up to 28% dcw in a continuous form in a photobioreactor (PBR) operational for 108 days [36]. EPS synthesis has been explored to a much lesser extent. Cyanobacteria microbiomes offer a unique ecological context where inter-strain interactions, cooperative behaviours, and synergistic metabolic activities may lead to enhanced biopolymers synthesis compared to individual strains in monoculture.

Expanding the horizons of bioprocessing through exploring coproduction of valuable metabolites can significantly improve the overall bioprocess efficiency by generating multiple products from a single cultivation. Previous research on production of both extracellular and intracellular compounds in this area is very limited. To our knowledge, studies with cyanobacteria have only investigated combinations such as EPS and phycobiliproteins [37–39], EPS and ethanol [40,41]; PHB and phycobiliproteins [42,43]. There is only one reference on PHB and EPS [44].

Recognizing the significance of this gap, our study aims to contribute to address this complexity by focusing on cyanobacteria microbiomes to explore the simultaneous PHB and EPS production. We employed seven cyanobacteria rich cultures collected from field environmental samples, that we previously tested for PHB production [35,36,45]. Our goal was to screen for potential EPS- producers and evaluate the effect of two parameters (salinity and presence of acetate as an external organic carbon source) on EPS synthesis. These two parameters were chosen based on their previous known effects either on EPS of PHB: salinity has been observed to impact EPS production, while Ac improves PHB production (which in turn could affect EPS). Though the effect of using organic carbon sources, like glucose, lactose, or sucrose, in EPS production in cyanobacteria has been reported [44,46,47], the influence of Ac on EPS production has received no attention until now. In this investigation, Ac was deliberately chosen as carbon source to investigate whether Ac also induces EPS production and explore the potential integration of EPS production with PHB synthesis. Finally, we scaled-up the process in a 3 L PBR to evaluate the feasibility of simultaneous production of EPS and PHB.

## Materials and Methods

### Inoculum

Seven microbiomes collected in [45] were used as inoculum. The sample code stablished in Altamira2023 for each sample is adopted in this text (Table 1).

**Table 1.**
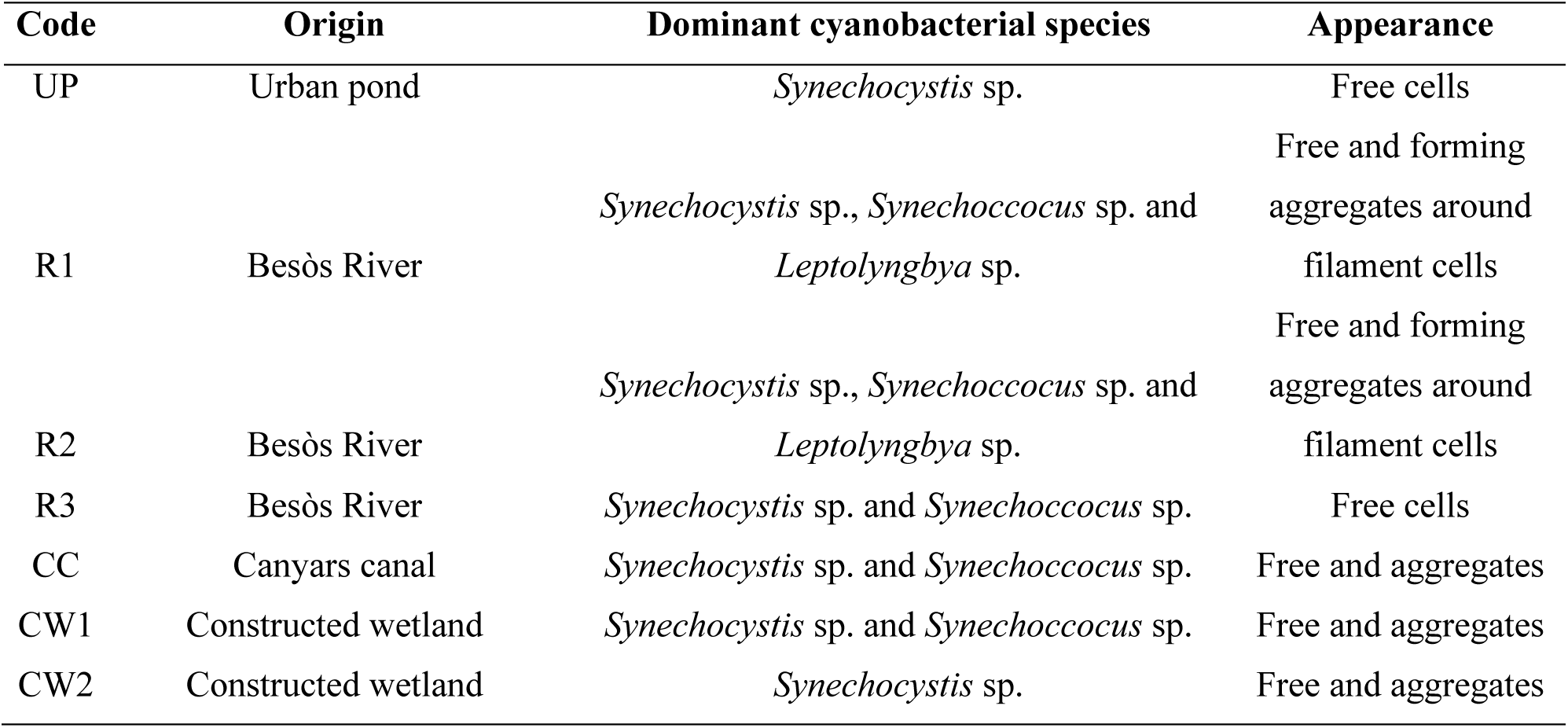
Sample code, origin of samples and microscopic appearance of the cultures used in this study.

Samples were collected from (i) an urban pond located in a park (Barcelona, Spain), (ii) Besòs river (Sant Adrìa de Besòs, Spain,), (iii) Canyars canal outlet close to the sea (Gavà, Spain) and (iv) the constructed wetland in Can Cabanyes (Granollers, Spain). They were kept in BG-11 medium with low phosphorus (P) concentration (0.1 mgP·L^-1^) to favour cyanobacteria growth over other phototrophs, such as green algae. Cyanobacterial species *Synechocystis* sp., *Synechoccocus* sp. and *Leptolyngbya* sp. dominated the cultures used as inoculum (Table 1 and Figure A1). Their presence and appearance were validated by bright light and fluorescence microscope observations (Nikon, Japan), and their taxonomy classification was prior conducted by 16S rRNA gene amplification [45].

### Experimental design

To determine the optimal cultivation conditions for EPS production by the seven microbiomes, response surface methodology (RSM) was applied [48]. This approach allowed to assess the impact and interactions of two key experimental variables (coded as X), (X_1_) salt stress and (X_2_) the presence of an external organic carbon source (acetate, Ac). To explore these responses comprehensively, a central composite rotatable design (CCRD) consisting of nine combinations was [49]. This design included four factorial design points at levels ±1 (trials 1-4), four experiments at axial level a = ±1.414 (trials 6-9), and a central point with three replicates (trial 5) (Table 2). The systems behaviour was evaluated by fitting the experimental data to a second-order polynomial model. The responses (Y) under investigation were (Y_1_) maximum EPS (including both RPS and CPS) concentration, and (Y_2_) the relative proportion of individual monomers, namely, fucose (Fuc), rhamnose (Rha), arabinose (Ara), glucosamine (GlcN), galactose (Gal), glucose (Glc), galacturonic acid (GalA) and glucuronic acid (GlcA).

**Table 2.** Design matrix of the CCRD applied to each microbiome with two independent variables (X_1_ and X_2_), NaCl and Ac concentration.

| | NaCl ($\text{g}\cdot\text{L}^{-1}$) | Acetate ($\text{g}\cdot\text{L}^{-1}$) |
| --- | --- | --- |
| Trial | ($X_1$) | ($X_2$) |
| 1 | 31 | 2.4 |
| 2 | 5 | 2.4 |
| 3 | 31 | 0.4 |
| 4 | 5 | 0.4 |
| 5* | 18 | 1.4 |
| 6 | 36 | 1.4 |
| 7 | 0 | 1.4 |
| 8 | 18 | 2.8 |
| 9 | 18 | 0 |

\*center point, done in three replicates.

To identify an appropriate reduced quadratic model, the significance of each source of variation was obtained from statistical analysis using software JMP®, version 14 (SAS Institute Inc.). A p- value< 0.05 was applied as the significant level.

### Experimental set up - 50 mL tubes test experiment

Conditions established by the experimental design were applied to the seven microbiomes. Inoculation was carried out in 50 mL Pyrex™ test tubes (Figure A2A) with approximately 1 g ·L-1 biomass (expressed as volatile suspended solids (VSS)). To prevent green microalgae competitors, 50 mL BG-11 medium as described in (rueda2020) with a modified P concentration (0.1 mgP·L⁻¹) was utilized. Sodium chloride (NaCl) and sodium acetate (NaAc) were added to the medium when necessary (Table 2). Tubes were continuously agitated via compressed air bubbling through a 0.22 µm pore filter and illuminated by cool-white LED lights, producing an intensity approx. 29 µmol·m-2 s-1 in a 15:9h light:dark photoperiod. The test lasted seven days and afterwards, EPS content was analysed as detailed below.

### Experimental set up - Simultaneous PHB and EPS production

Simultaneous PHB and EPS production was evaluated in a 3 L glass PBR (Figure A2B). Production was performed following the dual cycle approach described in [35]. Briefly, experiment started with a growth phase, where the PBR was inoculated with 100 mg VSS·L^-1^. BG-11 as described in [9] with modified concentrations of bicarbonate, as source of inorganic carbon (IC), nitrogen (N) and phosphorus (P), was used as media (100 mgIC L^-1^, 50 mgN·L^-1^ and 0.1 mgP·L^-1^). When N concentration was below 5 mg·L^-1^ (after 7 days), 600 mg Ac·L^-1^ were added and PBRs were enclosed during seven days with PVC tubes to avoid light penetration.

Reactors were continuously agitated by a magnetic stirrer ensuring a complete mixing and culture temperature was kept at 25-30 °C. During growth phase, illumination in the PBRs was maintained at 30 klx (approx. 420 µmol·m^-2^·s^-1^) using a 200 W LED floodlight positioned 15 cm from the reactors surface, operating in 15:9 h light:dark cycles. Throughout the growth phase, pH levels were regulated to remain within a range of approximately 8 to 8.3 using a pH controller system (HI 8711, HANNA instruments). This system activated an electrovalve to inject CO_2_ into the reactors when the pH reached 8.3, subsequently adjusting it back to 8. pH data was recorded at 5 min intervals using the PC400 software (Campbell Scientific). During the dark phase, the pH was measured but not actively controlled as photosynthesis was not occurring.

This experiment was done in triplicate and results are shown as the mean values ± standard deviation.

### Analytical methods

The Ac concentration in the medium at the end of the 50 mL tubes test experiment was analyzed following [50]. After centrifugation, the supernatant was filtered (0.2 µm membrane) and Ac concentration was determined by HPLC using a refractive index detector and a BioRad Aminex HPX-87H column. The analyses were performed at 50 °C, with 0.01 N H_2_SO_4_ as eluent at a flow rate of 0.6 mL·min^-1^. To normalize the results and compare Ac consumption across the different conditions and microbiomes tested, the following Equation (1) was employed:

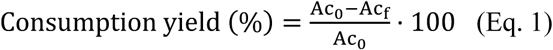

where Ac_f_ represents the acetate concentration at the end of the experiment (day 7) and Ac_0_ is the initial concentration of each trial.

During simultaneous production test of PHB and EPS, biomass concentration was determined as VSS according to procedure 2540-D described in Standard Methods [51]. Turbidity was measured with a turbidimeter (HI93703, HANNA Instruments). To provide a rapid estimation of biomass concentration, VSS were correlated with turbidity (Figure A3). To analyse the concentration of nitrogen and Ac in the PBRs, samples were collected and filtered through a 0.7 μm pore glass microfiber filter. Nitrogen analysis was conducted during the growth phase, following method 4500-NO^-^_3_ (B) from Standard Methods [51]. Note that in BG-11 the only source of N is nitrate. Ac was analysed with the acetate colorimetric assay kit MAK086 (Sigma-Aldrich) following supplier instructions. Samples were analysed with a BioTek Synergy HTX plate reader (Agilent Technologies) set to 450 nm.

### Glycogen extraction and quantification

Glycogen (Gly) analysis was done following the method described by [52] with minor modifications. In brief, freeze-dried biomass (2 mg) was mixed with 2 mL of 0.9 M HCl and subjected to digestion for 3 h at 100 °C. Sample was centrifuged (12,000 g for 2 min) and the supernatant was filtered (0.2 µm membrane). Finally, glucose was analysed by anion exchange chromatography, using a Metrosep Carb 2–250/4.0 column (Agilent Technologies), equipped with a pulsed amperometric detector. The eluent used was 300 mM sodium hydroxide and 1 mM sodium acetate. The analysis was performed at 30 ◦C, at a flow rate of 0.5 mL min^−1^. Glucose standard was used at concentrations in the range of 5 to 100 ppm.

### EPS extraction

After the seven days 50 mL tubes test experiment, the culture broth was centrifuged at 13,000 g for 20 min. The cell-free supernatant was used for the RPS extraction and quantification; and the cell pellet was used for CPS extraction and quantification.

The methodology described in [53] was used with some modifications. The culture broth was centrifuged at 8,000 g for 20 min. To extract RPS, the cell-free supernatant was submitted to a dialysis with 12-14 kDa MWCO membrane (Spectra/Por^®^, Spectrum Laboratories, Inc.) against deionized water, at room temperature, under continuous stirring. The dialysis water was changed frequently until conductivity of the water reached a value below 10 µS cm^−1^ (approximately after 48 h). Finally, samples were frozen at-80 °C and freeze-dried.

To extract CPS, the biomass pellet was rinsed with 2 mL saline buffer (2 mM Na_2_HPO_4_·2 H_2_O, 4 mM NaH_2_PO4·12H_2_O, 9 mM NaCl, 1 mM KCl, pH 7.0) following method 3 described in silva2017. Briefly, samples in saline buffer were sonicated with an ultrasound bath (Bandelic electronic GmbH & Co) for four cycles of 30 s, alternating with 30 s in ice. Samples were left overnight at-20 °C and the supernatant was recovered by centrifugation (12,000 g for 20 min at 4 °C). Then, the same methodology applied to extract RPS was used.

EPS volumetric production rate (ϒ_EPS_ (mgEPS·L^-1^·d^-1^)) was obtained by:

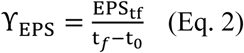

where EPS_*tf*_ is the concentration of EPS (in mg·L^-1^) quantified at the end of the experiment (day 14, t_f_).

The EPS yield on acetate (Ac) (*Y*_EPS/AC_) was calculated on a Chemical Oxygen Demand (COD)- basis by:

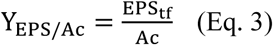

where EPS_*tf*_ is the concentration of EPS (given 1.36 gCOD·gEPS^-1^ [54]) end of the experiment (t_f_). Ac (mg·L^-1^) is the acetate concentration (given 1.07 gCOD·gAc·L^-1^) added (600 mgAc·L^-1^) in the medium at the beginning of the dark phase.

### EPS composition analysis

For the EPS compositional analysis, methodology described in [55] was followed. Freeze dried samples (1 mg) were dissolved in 1 mL deionized water and hydrolysed with trifluoroacetic acid (TFA) (0.02 mL TFA 99%) at 120 °C, for 2 h. The hydrolysate was used for the identification and quantification of the constituent monosaccharides.

Samples from the 50 mL tubes test experiment were analyzed by HPLC, using a CarboPac PA10 column (Dionex), equipped with pulsed amperometric detector. The analysis was performed at 30 °C, at an eluent flow rate of 1 mL min^-1^, with the following eluent gradient: 0–20 min, sodium hydroxide 18 mM; 20–40 min, sodium hydroxide (50 mM) and sodium acetate (170 mM). Fucose, rhamnose, arabinose, glucosamine, galactose, glucose, mannose, glucuronic acid and galacturonic acid at concentrations between 1 and 100 ppm were used as standards.

Samples from the simultaneous production test of PHB and EPS, were analysed by anion exchange chromatography, using a Metrosep Carb 2–250/4.0 column (Agilent Technologies), equipped with a pulsed amperometric detector. The eluents used were (A) 1 mM sodium hydroxide and 1 mM sodium acetate, and (B) 300 mM sodium hydroxide and 500 mM sodium acetate. The analysis was performed at 30 ◦C, at a flow rate of 0.6 mL min^−1^, with the subsequent eluent gradient: 0–22 min, eluent (A); 22–30 min, eluent (A) 50% and eluent (B) 50%; 30-40; 22–30 min, eluent (A) 50% and eluent (B) 50%; 40-46min, eluent (A); and 46-60 min eluent (A). Fucose, rhamnose, galactose, glucose, mannose, xylose, and glucuronic acid at concentrations between 1 and 100 ppm were used as standards.

### PHB extraction and quantification

PHB was analysed at selected time points following methodology described in [56]. Briefly, 50 mL samples were taken and centrifuged (3,000 g for 10 min). Cell pellet was frozen at-80°C overnight. The frozen samples were then freeze-dried for 24 h (−110 °C, 0.05 hPa). Freeze-dried biomass (3-3.5 mg) was mixed with 1 mL CH_3_OH with H_2_SO_4_ (20% v/v) and 1 mL CHCl_3_ containing 0.05 % w/w benzoic acid. The samples underwent heating for 5 h at 100 °C in a dry-heat thermo-block, followed by cooling in a cold-water bath for 30 min. Subsequently, 1 mL of deionized water was added, and the tubes were vortexed for 1 min. The CHCl_3_ phase, containing dissolved PHB, was recovered and introduced into a chromatography vial with molecular sieves. Gas chromatography analysis (7820A, Agilent Technologies) was performed using a DB-WAX 125-7062 column. Helium served as the gas carrier (4.5 mL·min-1), with an injector split ratio of 5:1 and a temperature of 230 °C. The flame ionization detector (FID) temperature was set to 300 °C. A standard curve of the co-polymer PHB-HV was utilized for PHB quantification. PHB volumetric production rate (ϒ_PHB_ (mgPHB·L^-1^·d^-1^)) was obtained by:

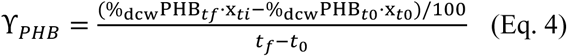

where %_dcw_PHB_*tf*_and %_dcw_PHB_*t*0_ are the yield on biomass quantified at the end of the experiment (t_f_) and at the beginning of the dark phase (t_0_). X_*tf*_ and X_*to*_ are the biomass concentration (in mgVSS·L^-1^) at the end and at the beginning of the dark phase, respectively.
The PHB yield on acetate (Ac) (*Y*_PHB/AC_) was calculated on a COD-basis by:

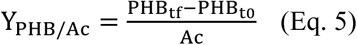

The amount of PHB produced (given 1.67 gCOD·gPHB^-1^) was obtained by multiplying the %dcw PHB produced per biomass concentration (in mgVSS·L^-1^) at the end of the test (t_f_) and at the beginning (t_0_) of the dark phase. Ac (mg·L^-1^) in the equation is the acetate concentration (given 1.07 gCOD·gAc·L^-1^) added (600 mgAc·L^-1^) in the medium at the beginning of the dark phase.

### EPS and PHB staining

By the end of the simultaneous production test for microbiomes R1, both bioproducts were visualized by staining analysis. A simple staining with black Chinese ink was conducted for EPS visualization, and observation was carried out under a bright light microscope (Nikon, Japan). For PHB staining, 1 % (wt/vol) Nile Blue A solution was used, with samples examined via Confocal Laser Scanning Microscope (CLSM), as detailed below.

### Confocal Laser Scanning Microscope

Confocal Laser Scanning Microscopy (CLSM) images displaying intracellular PHB were obtained at the end of the simultaneous production test for microbiomes R1. Firstly, 2 mL of culture were centrifuged (6,000 g for 4 min). Cell pellets were then rinsed three times with PBS (200 µL) and fixed with a solution (400 µL) of glutaraldehyde (2.5 % in PBS) for 15 min, followed by three additional washes in PBS. Finally, 1 % (wt/vol) Nile Blue A solution was used for PHB staining. Stained samples were observed with a 63 x 1.4 numerical aperture oil immersion objective lens, excited with a diode 561 nm, and were viewed in a Carl Zeiss LSM 800 (Zeiss).

## Results and discussion

### Effect of salinity and acetate on EPS production

The influence of salinity and acetate (Ac) on EPS synthesis was delved across diverse conditions, ranging from 0 to 31 g·L-1 NaCl and 0 to 2.8 g·L-1 Ac, respectively. Following seven days of exposure to these varied conditions, the EPS production of each microbiome under each condition was evaluated. The selection of these parameters and their respective values was based on several factors. Firstly, the effect of salt on EPS synthesis has been observed to vary depending on the cyanobacterial strain [5,20]. Therefore, by examining a broad range of salinity levels, we aimed to capture potential strain-dependent responses. While acetate is well-known as an inducer of PHB synthesis [10,16,17,45], its impact on EPS production has been relatively underexplored in the literature. Hence, we chose to include acetate as a parameter to investigate its potential influence on EPS synthesis, thus expanding the understanding of the factors governing biopolymer production in cyanobacteria microbiomes.

The synthesis of RPS and CPS showed variations from one microbiome to another (Table 3 and Figure A4). Generally, the production of RPS and CPS was comparable, except for cultures UP, CC and CW1 where CPS synthesis was relatively higher than that of RPS (Table 3). Similarly, [26] also reported that RPS was formed to a lesser extent than CPS in *Anabaena* sp. and *Nostoc* sp.

**Table 3.**
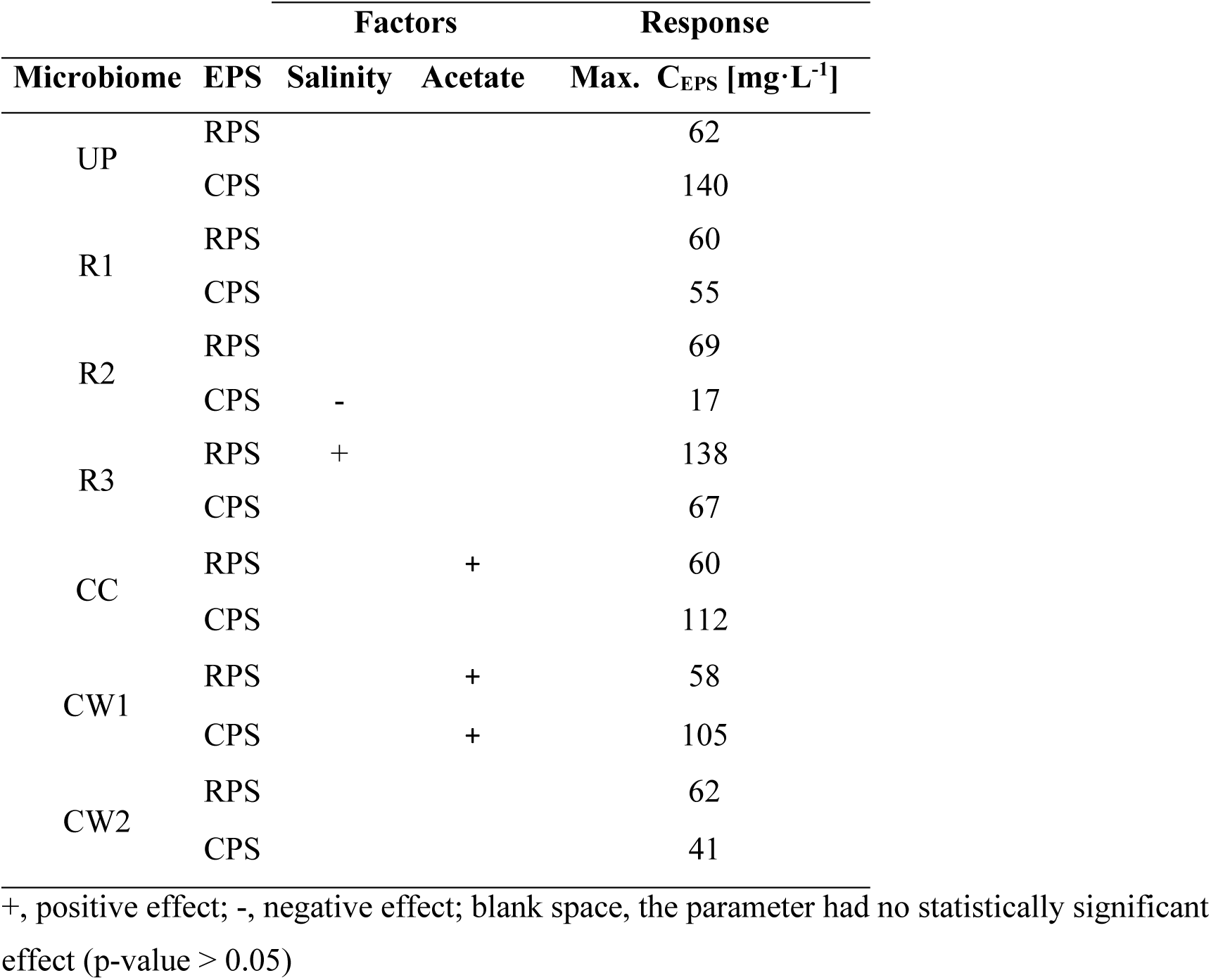
Summary of the statistically significant effects of the two factors evaluated (salinity and acetate) on EPS (RPS and CPS) synthesis in the seven microbiomes studied. The maximum EPS synthesis (C_EPS_) represents the highest concentration achieved by each microbiome across all conducted trials.

Among the seven cultures evaluated, microbiome R3 demonstrated the highest yield in RPS production, reaching a maximum of 138 mg·L^-1^ (Figure A4). This maximum was observed in trial 5, which involved the addition of 18 gNaCl·L^-1^ and 1.4 gAc·L^-1^. Prior to this, microbiome R3 had been tested for EPS production in another study [45], resulting in a modest synthesis of 7 mg·L^-1^ EPS with the addition of 0.04 gAc·L^-1^. It is noteworthy that Ac did not statistically affect EPS production (p > 0.05); however, the reasons behind this lack of statistical significance were not explained by the authors. Interestingly, our recent findings align with these earlier observations; similarly, Ac did not significantly impact EPS production, suggesting that the considerable increase in polymer output can be attributed to the presence of salt. This indicates salt’s positive influence on RPS production specifically in this culture (Table 3). Notably, microbiome R3 featured a higher abundance of *Synechococcus* sp. within its community (Figure A1). This prevalence of this cyanobacteria might account for the observed enhancement in RPS production in response to salt addition. Similarly, this positive correlation between NaCl exposure and EPS production has also been documented in other cyanobacteria, including *Synechocystis* sp., *Nostoc* sp. and *Spirulina* sp. [28,57,58]. Particularly in the case of *Nostoc* sp., CPS synthesis was higher (212 mg·L^-1^) under salt stress compared to the control group (126 mg·L^-1^) [57]. However, it is noteworthy that salt stress typically leads to a simultaneous decrease in biomass growth. High concentrations of salts, such as NaCl, can disrupt the osmotic balance inside and outside the cells, causing water loss and potentially leading to cell lysis. This osmotic stress can impair the normal metabolic processes and growth rates of cyanobacteria, affecting their overall biomass production [28,57]. Some cyanobacteria have developed mechanisms to tolerate and withstand high salinity levels by the stimulation of EPS synthesis that retain moisture around the cell preventing them of desiccation [59,60]. The ability of cyanobacteria to respond to high salinity can vary greatly among different species. Indeed, NaCl did not induce a significant change in EPS synthesis in cultures of *Spirulina* sp, *Anabaena* sp., *Synechocystis* sp. or *Cyanothece* sp. [23,25,26,61]. [61] investigated the effects of high NaCl (17.5 g·L^-1^) and low sulphur concentration (1.2 mg·L^-1^) on RPS production in *Synechocystis* sp. Surprisingly, neither condition led to a significant increase or decrease in RPS production compared to the control group, where the concentration was below 100 mg·L^-1^ in all scenarios.

Differences were noted among the evaluated microbiomes regarding Ac consumption (Table A1). Notably, only cultures CC and CW1, which demonstrated the highest rated of Ac consumption, experienced a beneficial impact on EPS synthesis following the addition of Ac to the medium (Table 3). In fact, a strong correlation between Ac consumption and EPS production was evident, with a correlation coefficient (R^2^) higher than 0.7 in each culture (Figure A5). Specifically, culture CW1 saw enhancements in both RPS and CPS synthesis, whereas culture CC experienced a boost primarily in RPS synthesis (Table 3). Noteworthy is the substantial increase observed in culture CW1, particularly in trials 1, 2, 5-8, where the presence of Ac led to a substantial enhancement. Similarly, in culture CC, the final CPS concentration rose up to six times compared to trials without Ac (Figure A4), underscoring the impact of Ac on EPS synthesis in these cultures, due to Ac consumption. This observation was consistent with previous findings where Ac supplementation significantly enhanced EPS production in these both cultures [45]. Likewise, in *Nostoc* sp., presence of Ac boosted EPS synthesis [6,47], reaching levels around twice as high as those observed in autotrophic cultivation [47]. Nonetheless, it is worth noting that addition of acetate, valerate, glucose or citrate to the growth medium of *Anabaena* sp. led to a reduction in EPS concentration, although the authors did not provide an explanation for this phenomenon [44].

Furthermore, variations in the cyanobacterial species present in each microbiome, along with their distinct morphologies observed under the microscope (Table 1), did not result in noticeable differences in EPS production (Table 3). In fact, results on EPS (RPS and CPS) production by the evaluated microbiomes were similar and comparable to that obtained in monocultures of various cyanobacteria [39,46,62–65]; however, higher production has also been documented in cultures of *Nostoc* sp. and *Anabaena* sp., with synthesis reaching over 1,000 mg·L^-1^ in various days at stationary phase [6,63]. These findings underscore the complexity of microbial responses to environmental stimuli and highlight the need for further research to unveil the underlying mechanisms governing biopolymer production in microbial communities. Nevertheless, moving forward, an essential consideration is the adaptation of culture and operational conditions to obtain polysaccharides with desired properties tailored to their intended application. These properties will be linked to the composition of the EPS.

### Effect of salinity and acetate on EPS monosaccharides composition

Analysing the monomer composition of EPS is crucial in comprehending their functional significance. To accurately reflect the EPS sugar composition of each, the average RPS and CPS composition for each culture has been calculated based on the results obtained in each trial for every microbiome (Table 4).

**Table 4.** Monosaccharide and non-sugars groups’ composition (in relative proportion, %) of EPS produced by the microbiomes tested in this study. Values are the average and standard deviation obtained under conditions described in Table 2.

| Microbiome | EPS | Fuc | Rha | Ara | GlcN | Gal | Glc | Man | GalA | GlcA |
| --- | --- | --- | --- | --- | --- | --- | --- | --- | --- | --- |
| UP | RPS | 2.7 ± 0.6 | 6.4 ± 1.9 | 1.4 ± 0.3 | 6.0 ± 1.4 | 4.6 ± 1.2 | 44.3 ± 8.5 | 34.1 ± 5.2 | 0.7 ± 0.2 | 0.7 ± 0.2 |
|  | CPS | 1.0 ± 0.8 | 11.6 ± 3.3 | n.d. | n.m. | 6.6 ± 8.3 | 29.1 ± 18.4 | 56.9 ± 16.9 | 7.4 ± 2.3 | 3.7 ± 1.3 |
| R1 | RPS | 3.0 ± 1.8 | 4.9 ± 0.3 | 1.0 ± 0.3 | 5.7 ± 2.0 | 7.2 ± 1.4 | 47.4 ± 11.7 | 32.7 ± 7.1 | n.d. | 0.8 ± 0.2 |
|  | CPS | 4.7 ± 3.4 | 11.8 ± 4.7 | 1.1 ± 0 | n.m. | 2.7 ± 1.3 | 52.4 ± 14.3 | 32.8 ± 15.2 | 1.7 ± 0.3 | 2.4 ± 1.6 |
| R2 | RPS | 2.4 ± 0.8 | 2.9 ± 1.3 | 1.0 ± 0.4 | 4.7 ± 2.1 | 6.8 ± 2.0 | 48.9 ± 11.6 | 30.0 ± 6.3 | n.d. | 1.0 ± 0.6 |
|  | CPS | 11.1 ± 0 | 16.9 ± 11.4 | 2.7 ± 0 | n.m. | 5.8 ± 4.9 | 43.5 ± 15.42 | 57.2 ± 24.4 | n.d. | 7.3 ± 11.5 |
| R3 | RPS | 6.6 ± 2.8 | 11.3 ± 15.3 | 2.6 ± 0.9 | 3.7 ± 1.2 | 5.0 ± 1.8 | 50.6 ± 18.3 | 19.6 ± 7.5 | n.d. | 2.1 ± 0.9 |
|  | CPS | 9.5 ± 3.3 | 16.8 ± 3.8 | 2.3 ± 1.9 | n.m. | 3.0 ± 1.6 | 44.3 ± 19.7 | 24.5 ± 4.7 | 1.4 ± 0.4 | 2.7 ± 3.9 |
| CC | RPS | 5.2 ± 1.6 | 8.4 ± 3.2 | 1.7 ± 0.4 | 8.3 ± 1.5 | 5.9 ± 0.8 | 38.6 ± 4.6 | 28.7 ± 3.4 | 1.7 ± 0.7 | 1.8 ± 0.6 |
|  | CPS | 1.9 ± 1.6 | 20.3 ± 8.4 | n.d. | n.m. | 2.0 ± 1.5 | 18.9 ± 8 | 57.2 ± 14.8 | n.d. | 4.4 ± 3.0 |
| CW1 | RPS | 4.0 ± 1.6 | 6.3 ± 6.3 | 1.4 ± 1.2 | 6.6 ± 2.3 | 7.6 ± 2.1 | 51.5 ± 11.0 | 23.5 ± 6.7 | n.d. | 1.0 ± 0.5 |
|  | CPS | 8.1 ± 2.3 | 16.1 ± 5.1 | 3.0 ± 1.7 | n.m. | 3.2 ± 1.4 | 32.6 ± 5.9 | 35.3 ± 5.6 | n.d. | 5.2 ± 5.0 |
| CW2 | RPS | 4.4 ± 1.0 | 7.9 ± 3.4 | 1.7 ± 0.5 | 6.7 ± 1.6 | 5.3 ± 0.9 | 36.4 ± 7.7 | 39.3 ± 4.3 | n.d. | 0.8 ± 0.4 |
|  | CPS | 4.8 ± 2.1 | 14.0 ± 3.2 | 1.7 ± 0.7 | n.m. | 2.9 ± 1.1 | 23.5 ± 13.6 | 52.3 ± 19.2 | 5.3 ± 1.7 | 7.3 ± 8.7 |

\*Ara, arabinose; Fuc, Fucose; Gal, galactose; GalA, galacturonic acid; Glc, glucose; GlcA, glucuronic acid; GlcN, glucosamine; Man, mannose; Rha, rhamnose; n.d., non-detected; n.m., non-measured

Cyanobacterial EPS primarily consists of neutral sugars, including glucose, mannose, galactose, rhamnose, arabinose and fucose. Notably, our analysis revealed differences in monosaccharide composition between RPS and CPS (Table 4), aligning with the known complexity of cyanobacterial polymers [21,64,66]. Generally, glucose is the main monosaccharide in cyanobacteria EPS [5,26,28,57,67,68]. Consistent with these findings, in both RPS and CPS, glucose, together with mannose, emerged as the most common monosaccharides across the seven microbiomes tested, making up around 60-80 % of the total polysaccharide. The predominance of glucose and mannose were consistent with that observed for EPS produced individually by *Anabaena* sp., *Microcystis* sp. *Cyanothece* sp., *Nostoc* sp., *Synechocystis* sp. or *Synechococcus* sp.; although ratios may differ between species [28,57,66,67,69–72]. Additional monosaccharides, including galactose, arabinose, rhamnose and fucose, were present in the RPS from the seven evaluated microbiomes (Table 4). Although their ratio was relatively low (< 11 % of the total RPS), values agree with that reported in other cyanobacteria [24,26,66,70–72]. Notably, glucosamine was present in all RPS, comprising an average proportion of 6% of the total RPS, aligning with previous studies with cyanobacteria *Anabaena* sp., in which similar molar ratios were detected in RPS (4.7 %) and CPS (5.9 %) [26]. In CPS, proportion of rhamnose was around 11-20 % of the total CPS, while in RPS proportion of rhamnose was lower than 11 % (Table 4). Similarly, in cultures of *Anabaena* sp., major proportion of rhamnose was detected in CPS than in RPS [26].

The sugar composition remained consistent despite changes in salinity levels or the addition of Ac to the culture media. This finding is consistent with studies on EPS from *Synechocystis* sp., *Aphanothece* sp., *Microcystis* sp. and *Nostoc* sp., where the profile of monosaccharide composition remained unchanged despite variations in carbon sources or salinity [6,28,57,59,73]. However, a unique observation was made in the case of microbiome R3, where exposure to NaCl led to a significant alteration in the sugar composition of RPS. This shift mirrors prior findings in the relative sugar content of RPS from *Synechocystis* sp., where high NaCl (17.5 g·L^-1^) exposure resulted in higher levels of glucose, as well as, lower levels of galactosamine [61]. In EPS produced by *Nostoc* sp., the molar ratio of sugar residues underwent slight modifications with Ac supplementation, being the most significant change the decrease in mannose (50 % molar ratio versus 30 % without Ac) [6]. Other variations, including presence of nitrogen, have minimal effect on the monosaccharide composition and morphology of EPS [65]. Interestingly, despite significant variations in sugar ratios and potential morphological adjustments in both RPS and CPS from *Nostoc* sp. under different light wavelengths, the core structural elements of EPS appear to maintain their integrity, as evidenced by Fourier transform infrared spectroscopy and X-ray diffraction characterizations [72]. This observation suggests the existence of a highly regulated biosynthesis pathway that preserves essential structural characteristics of these polymers under different environmental conditions.

Uronic acids, identified as glucuronic and galacturonic acids, are also characteristic components of microalgae and cyanobacteria EPS. Sometimes in relatively high levels, like in *Phormidium* sp*., Neorhodella* sp. or *Heterocapsa* sp, where they have shown to represent around 30 % of the overall composition [74,75]. However, in other microalgae and cyanobacteria this ratio is lower, not higher than 17 % in *Anabaena* sp., 10 % in *Microcystis* sp., and even lower or not detected in species including *Cyanothece* sp., *Synechocystis* sp., *Nostoc* sp., *Synechoccocus* sp. or *Parachlorella* sp. [28,57,66,67,69–71]. Congruent with most results, uronic acids were almost undetected in RPS of the seven evaluated cultures (Table 4), where proportion did not exceed 2 % of the total polysaccharide. Interestingly, in CPS, their proportion was higher in all the samples, but still the average ratio was not more than 7 % of the molar ratio (Table 4).

These differences in RPS and CPS composition appear to be a question of discussion in the literature [21,65,70,73,76], and these variations were also evident in the present study. This is attributed to the distinct roles of these polysaccharides, where each component conveys unique characteristics. Remarkably, the relatively high glucosamine ratio (Table 4) suggest that these microbiomes hold potential as sources of novel and improved biomaterials for health applications [53]. Deoxysugars like rhamnose and fucose contribute for hydrophobic properties, while uronic acids contribute to the anionic and sticky nature of the polysaccharides [77]. This distinction was particularly pronounced in the CPS of the seven microbiomes, where these components were present in a higher proportion than in RPS (Table 4). Indeed, the presence of a polysaccharidic layer enveloping the cells suggests a potential mechanism to prevent direct contact between cells and toxic heavy metals present in the environment [76,78].

### Large photobioreactor cultivation – Simultaneous PHB and EPS production and characterization

The aim of the 50 mL test tubes experiment was to evaluate the EPS-producing abilities of seven microbiomes and examine the impact of salt and acetate on polysaccharide synthesis. This pursued to gain understanding of EPS synthesis by these cultures, ultimately leading to the integration of EPS synthesis with PHB production. Since neither salt or Ac improved considerably EPS production, we assessed this dual production under conditions optimized for PHB synthesis [35,36]. To accomplish this, microbiome R1 was cultivated in a 3 L PBR following a dual phase approach. The methodology involved a biomass growth phase followed by a dark incubation period where Ac is added into the medium upon N limitation.

Initially, microbiome R1 was inoculated into the 3 L PBR with a biomass concentration of 100 mg VSS·L^-1^. The biomass growth phase lasted seven days, in which the concentration reached an average 750 mgVSS·L^-1^ (Figure 1A). At this time point, the concentration of N was 5 mg·L^-1^ for all the trials performed Figure 1A). Then, the PBR was enclosed with PVC tubes to avoid light penetration and 600 mg·L^-1^ of Ac were added to the reactor.

**Figure 1.**
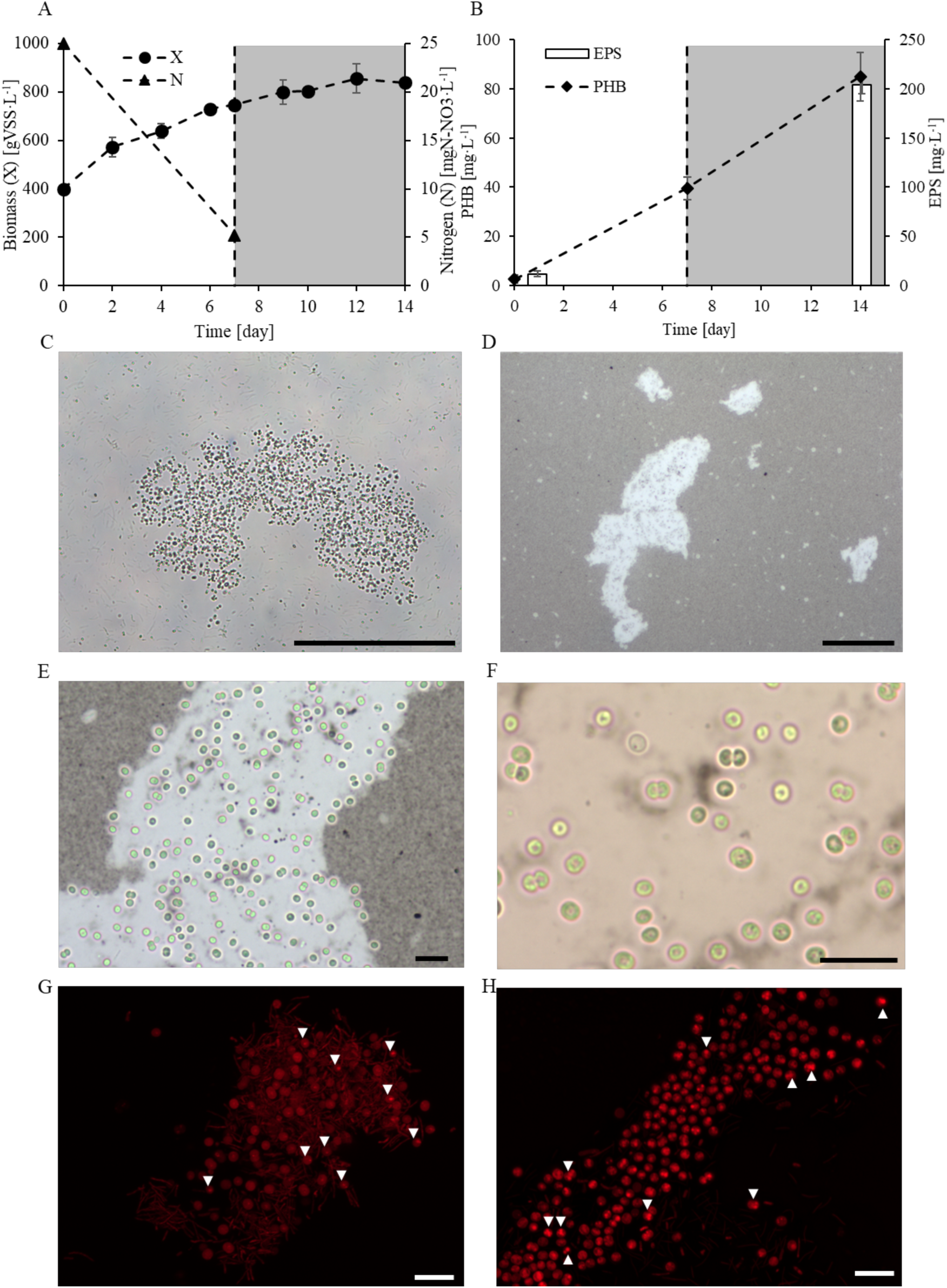
Concentration dynamics in (A) biomass (as VSS) and nitrogen (as N- NO_3_); and (B) PHB and EPS for microbiome R1 through the experiment. White and grey colour in the figure indicates growth phase and dark phase, respectively. Dashed line remarks the beginning of dark phase. Note that nitrate was not measured in the dark phase. These data represent the mean ± std of three experiments performed. (C) Aggregate of *Syenchocystis* sp. and *Syenchoccous* sp. in bright light microscope of microbiome R1 in 10X. Scale bar is 100µm. (D) Bright light microscope images of microbiome R1 after black Chinese ink staining at 10X; (E) at 40x, and (F) at 100X. EPS appear as light and shiny around the cells due to ink staining. Scale bar is 100µm in (D); and 10 µm in (E) and (F). (G) and (H) LCSM image of an aggregate of microbiome R1. White arrowheads point PHB granules within the cells. Scale bar is 10µm.

Interestingly, regarding EPS synthesis, both RPS and CPS values surpassed those obtained in our prior experimental design with the 50 mL tubes (Table 5). This suggests that the methodology employed may offer potential advantages for EPS synthesis. Alternatively, the increase could be attributed to the extended duration of the assay. Notably, by the end of the experiment, microbiome R1 achieved RPS concentrations of 77 mg·L^-1^ and CPS concentrations of 128 mg·L^-1^. Consistent with observations from the preceding experimental setup (Table 3), R1 demonstrated a propensity to produce higher CPS compared to RPS.

**Table 5.** EPS (RPS and CPS) and PHB production (in terms of mg·L^-1^ and %dcw). Values were calculated at the end of the experiment; the mean value obtained from the three tests conducted is shown together with the standard deviation.

| **Parameter** |  |
| --- | --- |
| **RPS (mg·L<sup>-1</sup>)** | 76.8 ± 7.5 |
| **CPS (mg·L<sup>-1</sup>)** | 127.5 ± 1.6 |
| **PHB (mg·L<sup>-1</sup>)** | 96.3 ± 28.7 |
| **PHB (%dcw)** | 11.5 ± 3.5 |
| **Y<sub>EPS</sub> (mgEPS·L<sup>-1</sup>·d<sup>-1</sup>)** | 14.6 ± 0.7 |
| **Y<sub>PHB</sub> (mgPHB·L<sup>-1</sup>·d<sup>-1</sup>)** | 5.6 ± 1.1 |
| **Y<sub>EPS/Ac</sub> (g EPS<sub>COD</sub> · g Ac<sub>COD</sub><sup>-1</sup>)** | 0.4 ± 0.0 |
| **Y<sub>PHB/Ac</sub> (g PHB<sub>COD</sub> · g Ac<sub>COD</sub><sup>-1</sup>)** | 0.1 ± 0.0 |

The monosaccharide composition remained consistent with previous analyses (Table 4), with glucose identified as the primary component in both RPS and CPS, comprising over 60 % of the polysaccharide (Table 6). Notably, there was a reduction in the ratio of mannose to below 10 %, contrasting with the 30 % ratio found in the previous test. Additionally, slight increases were noted in the proportions of fucose and rhamnose, reaching up to 8 % and 12 % of the total ratio, respectively. Furthermore, the sugar composition between RPS and CPS displayed similarities, with the ratios of monosaccharides being comparable across both types of polysaccharides. A notable exception was the presence of glucuronic acid, which was found in greater abundance in CPS compared to RPS, constituting 8% versus 3% of the total polymer, respectively.

**Table 6.** Monosaccharide and non-sugars groups composition (in relative proportion, %) of EPS (RPS and CPS) produced by microbiome R1 under simultaneously production of PHB. Note that in this case arabinose, glucosamine and galacturonic acid were not measured.

| **EPS** | **Fuc** | **Rha** | **Gal** | **Glc** | **Man** | **Xyl** | **GlcA** |
| --- | --- | --- | --- | --- | --- | --- | --- |
| RPS | 7.5 ± 1.3 | 12.1 ± 3.4 | 0.43 ± 0.1 | 62.4 ± 8.4 | 7.4 ± 1.5 | 7.7 ± 1.9 | 2.6 ± 0.9 |
| CPS | 6.0 ± 0.6 | 9.3 ± 2.6 | 0.51 ± 0.1 | 70.7 ± 16.2 | 5.7 ± 0.8 | 11.2 ± 09 | 8.2 ± 0.6 |

\*Fuc, Fucose; Gal, galactose; Glc, glucose; GlcA, glucuronic acid; Man, mannose; Rha, rhamnose; Xyl, xylose; n.d., non-detected

In relation to PHB synthesis, microbiome R1 reached an average 87 mgPHB·L^-1^ (9 % dry cell weight, dcw) by the end of the experiments performed (Table 5 and Figure 1B). These values were relatively lower than those reported for other cyanobacteria species operating under similar conditions (with Ac added at the initiation of dark incubation). For example, *Synechocystis* sp. demonstrated PHB accumulation of up to 22 %dcw by day 5 of incubation [17], *Chlorogloea* sp. reached up to 29 %dcw by day 6 [12], or a microbiome dominated by *Synechocystis* sp. accumulated up to 27 %dcw by day 7 [36]. The observed lower PHB accumulation values could be attributed to EPS synthesis (Figure 1B). Both PHB and EPS production are boosted by extracellular carbon sources, leading to competition for available exogenous carbon sources, like Ac [79–81]. Previous research on simultaneous PHB and EPS production by the bacteria *Sphingomonas* sp., the archaeon *Haloferax* sp., and microbial mixed cultures, has highlighted the impact of C/N ratio on biopolymer synthesis. A lower C/N ratio is often necessary for EPS synthesis, whereas a higher ratio is needed for maximum PHB accumulation [80,82,83]. Here, a high C/N ratio was employed, given that N was nearly depleted from the media, and 600 mg·L^-1^ Ac were added to the culture. Hence, it was expected that a higher PHB production should have been observed.

Despite these conditions suggesting increased PHB production, relatively low yield Y_PHB/Ac_ was obtained (Table 5) because acetate assimilation did not occur, as evidenced by the residual Ac concentration of 470 ± 100 mg·L^-1^ by the end of the tests. This clearly hindered PHB synthesis. In fact, a theoretical maximum yield Y_PHB/Ac_ could be achieved if all carbon from acetate was directed towards PHB synthesis. This would yield a maximum PHB concentration of 384 mgPHB·L^-1^, representing 52 %dcw PHB (considering 750 mgVSS·L^-1^, the average biomass concentration at the beginning of the dark phase). To enhance biopolymer production, exposing cells to alternating periods of growth and darkness shows potential for increasing PHB synthesis. This method enables cells to better adapt to conditions conducive to PHB synthesis [35,36], contrasting with the single-cycle experiments performed in this study.

Cyanobacteria also produce glycogen (Gly) as storage compound even without N or P limitation [84,85]. Microbiome R1 synthetized 130 mg·L^-1^ Gly during the growth phase, with concentration reaching an average 260 mg·L^-1^ (35 %dcw) by day 7 (Figure A6). In both mono-and mixed cultures of *Synechocystis* sp. and *Synechococcus* sp., similar Gly levels (20 – 30 %dcw) were detected following a growth period of 23 days [9] as well as shorter durations of 12 and 18 days [9,12]. Under the prolonged stress conditions in which cultures were submitted, characterized by nutrient deprivation and darkness during seven days, cells may have utilized the stored Gly as carbon reserve, potentially converting it into PHB [9,14,36,86,87]. However, there was no observed decrease in Gly content in the microbiome; rather, it remained stable (Figure A6). This stability correlates with the modest levels of PHB observed in the culture, suggesting that glycogen was not undergoing significant conversion into PHB. This phenomenon may be linked to EPS synthesis, using the IC added at the beginning of the test (100 mg·L^-1^), and redirecting carbon flux away from the glycogen metabolic pathways. Similarly, the disruption of *glgC* (encoding glucose-1-phosphate adenylyltransferase, involved in glycogen synthesis) in cyanobacteria *Synechocystis* sp. and *Synechococcus* sp. increased the EPS content by increasing the glucose concentration [40,68].

It seems evident that metabolic pathways associated with PHB were not highly active. As a response to the prolonged stress experienced by the cells, EPS production was triggered. Indeed, microscope observation revealed the presence of a mucilaginous external matrix enveloping the cells Figure 1C-F). Specifically, Chinese ink staining highlighted the EPS, conferring a light and shiny appearance to the matrix surrounding the biomass. This matrix formed an aggregate containing diverse cells, including unicellular cyanobacteria *Synechocystis* sp. and *Synechoccus* sp., as well as filaments in some of the aggregates (Figure 1 and A7). In microbiomes, the EPS layer plays a vital role in sustaining high-density populations of microorganisms and is a key factor in microbial flocculation [88,89]. This phenomenon is largely attributed to the presence of uronic acids in the polymer [59,63], accounting for 8 % of the CPS composition (Table 6). As cellular aggregation occurs, it accelerates the separation of biomass from the liquid medium, resulting in heightened efficiency. To visually appreciate this process, refer to Supplementary Video 1 for a time-lapse video.

In the video, the flask containing microbiome R1, which produces EPS, the presence of this polysaccharide significantly promotes biomass aggregation. This is evident from the photograph showing the culture stained with black Chinese ink, where the EPS appears as a white gel surrounding the cells. This visualization effectively demonstrates how EPS binds cells together, thereby accelerating the sedimentation process. The aggregated biomass forms a visible layer at the bottom of the flask. Conversely, in the flask with microbiome CW2, which does not produce EPS, the lack of such a substance, results in less effective biomass aggregation. Without EPS to facilitate cell-cell interactions and aggregation, the sedimentation rate is slower, and the culture broth remains more dispersed. This difference in culture broth appearance underscores the critical role of EPS in enhancing cellular aggregation and, by extension, the efficiency of the biomass aggregation process. Such improved efficiency in biomass separation would lead to notable cost savings in operations. Therefore, simultaneous production of PHB and EPS by cyanobacteria microbiomes presents a promising operational strategy, emphasizing practical benefits beyond the pursuit of high-value EPS production alone.

The presence of PHB was observed through polymer staining and visualization using LCSM. Nevertheless, PHB granules were clearly visualized as brightly fluorescent red granules within the cells (Figure 1D). Remarkably, cells consisted of a heterogeneous population with respect to PHB accumulation, as not all cells exhibited these granules (Figure 1D). This heterogeneous biopolymer content in the culture has been reported in other cultures due to the stochastic regulation of PHB synthesis [36,90–92].

## Conclusions

In conclusion, our study offers a comprehensive exploration of the EPS-producing capabilities within seven microbiomes enriched with cyanobacteria, mainly *Synechocystis* sp. and *Synechoccocus* sp. We investigated the influence of acetate and salt on polysaccharide production and sugar composition. While acetate supplementation or salt exposure did not yield significant alterations in overall EPS synthesis or composition, the positive response observed in two microbiomes (CC and CW1) to acetate supplementation was related to their higher Ac consumption in comparison to the other microbiomes. Conversely, salt exposure led to a statistically significant decrease in EPS synthesis in one microbiome (culture R2), while it positively influenced RPS production in R3. Despite these intricacies, a consistent EPS synthesis was observed across all tested conditions, with levels ranging from 25 to 150 mg·L^-1^, aligning with findings from similar cyanobacterial monocultures. Although differences were observed in the monosaccharide composition of RPS and CPS, both were identified as complex heteropolysaccharides. They were composed of six different monosaccharides, with glucose and mannose emerging as the predominant sugars among the EPS of the seven microbiomes studied. Together, these sugars constituted approximately 60 - 80 % of the total polysaccharide content.

Furthermore, our investigation into simultaneous EPS and PHB production in a 3 L PBR setup revealed challenges due to substrate competition. Although biopolymer synthesis was modest, the presence of uronic acid in the EPS facilitated biomass flocculation, streamlining the separation process, and potentially reducing associated time and costs. Finally, Nile Blue A staining revealed the internal PHB granules within cyanobacterial cells, while the use of black Chinese ink facilitated the visualization of the capsular EPS surrounding the cells.

Looking forward, refining strategies to regulate EPS production holds promise for enhancing the flocculation ability of PHB-producing cyanobacteria microbiomes. This represents a compelling avenue for advancing biopolymer production processes.

## CRediT authorship contribution statement

**Beatriz Altamira-Algarra:** Conceptualization, Validation, Formal analysis, Investigation, Writing – original draft. **Joan Garcia:** Conceptualization, Writing – review & editing, Supervision, Project administration, Funding acquisition. **Cristiana Torres:** Conceptualization, Supervision, Writing – review & editing**. Maria A. M. Reis:** Project administration, Funding acquisition. **Eva Gonzalez-Flo:** Conceptualization, Supervision, Writing – review & editing.

## Declaration of Competing Interest

The authors declare that they have no known competing financial interests or personal relationships that could have appeared to influence the work reported in this paper.

**Appendix A. Supporting information Supplementary**

