## Appendix A. Supporting information Supplementary for "Exploring simultaneous production of polyhydroxybutyrate and exopolysaccharides in cyanobacteria-rich microbiomes"

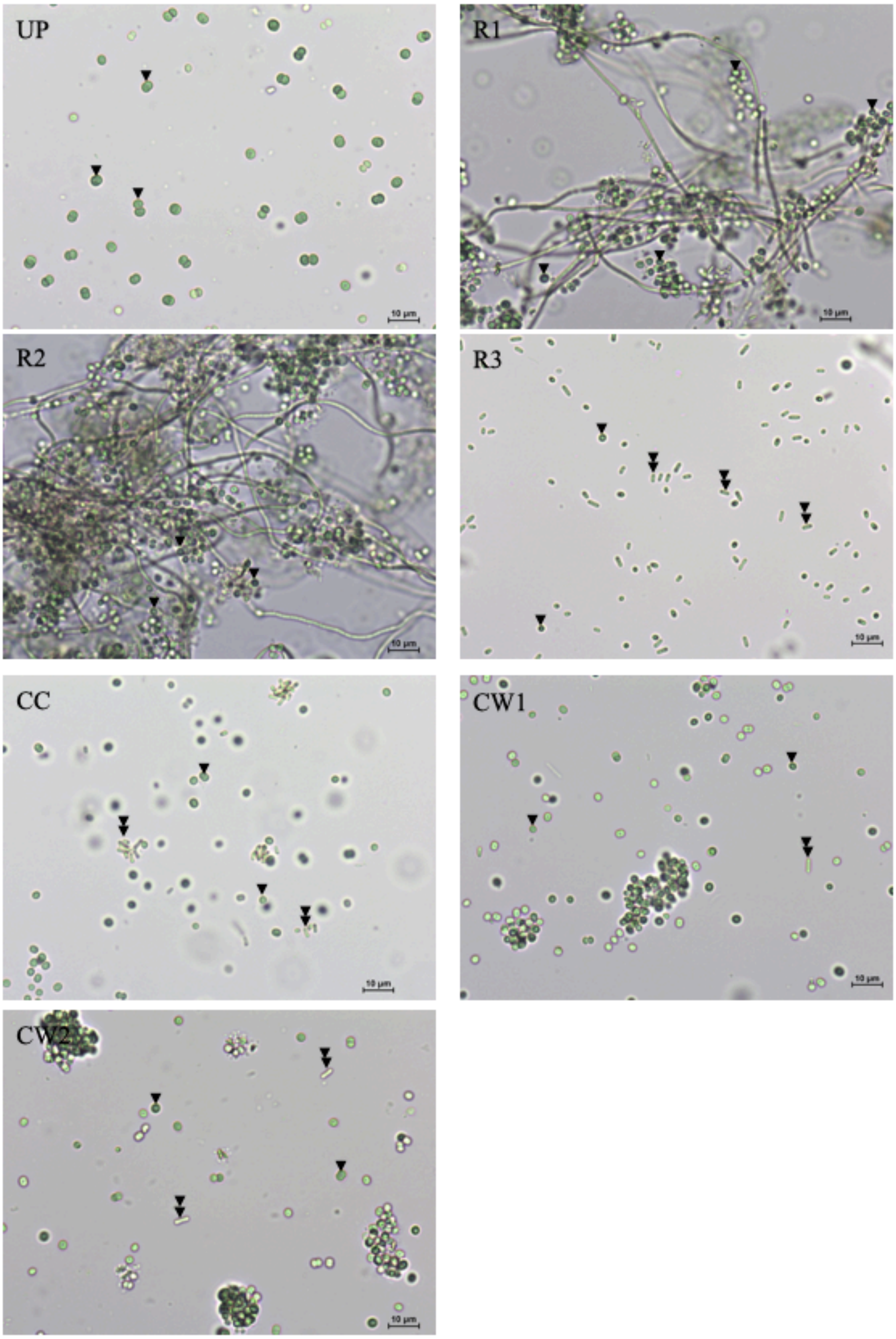


**Figure A1.** Bright light microscope images (40X) of microbiomes at start of the 50mL test tube experiment. Cyanobacterial species *Synechocystis* sp. and *Synechoccocus* sp. dominated the cultures. They were found as free cells or forming aggregates. Arrowhead points *Synechocystis* sp. and double arrowhead points *Synechococcus* sp. cells. Scale bas is 10 µm.

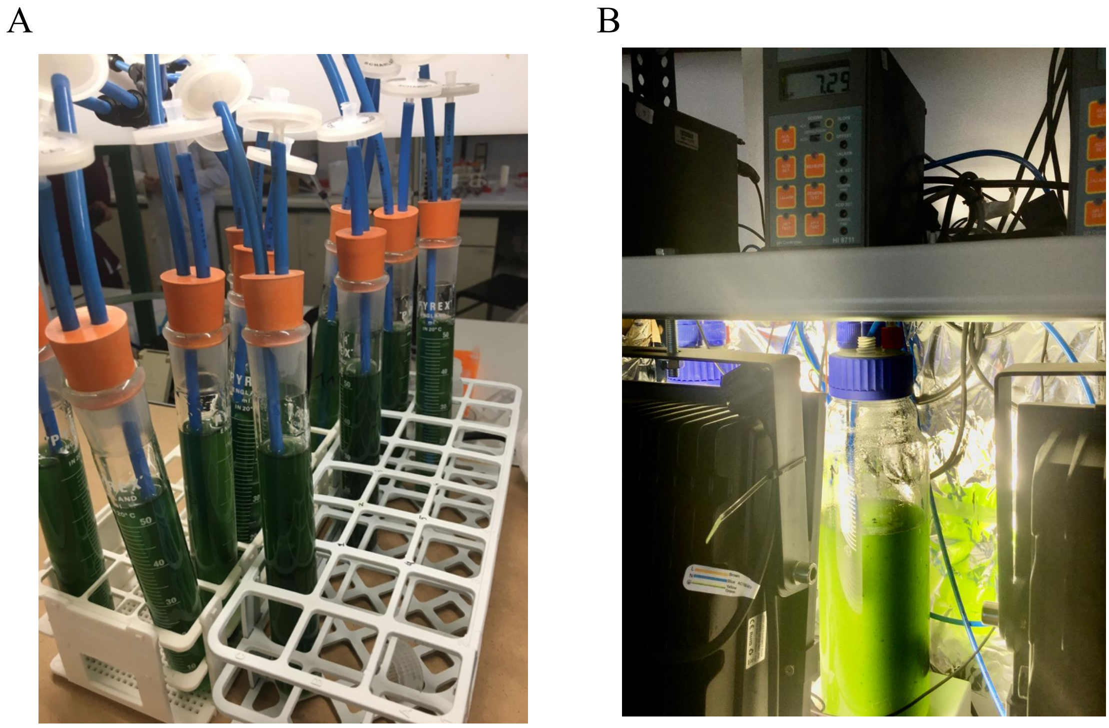


**Figure A2.** Image of the experimental set-up. (A) 50 mL tubes test experiment; and (B) 3 L PBR.

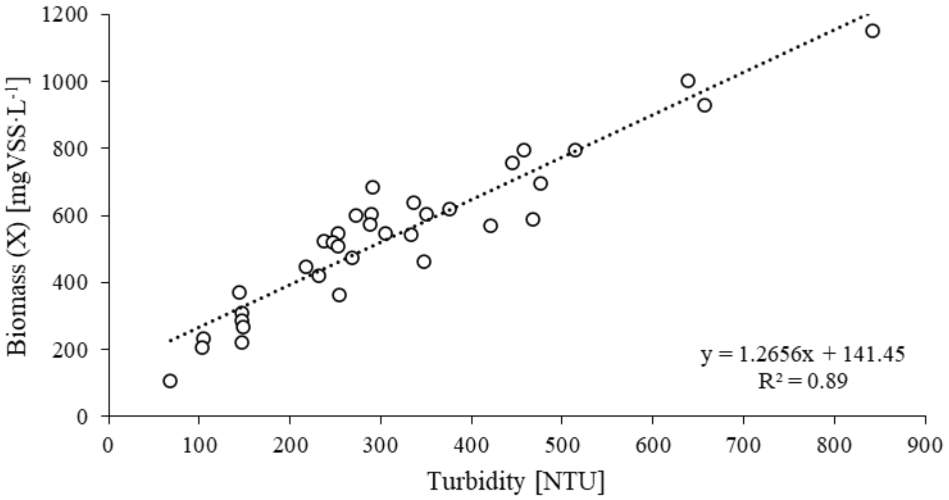


**Figure A3.** Calibration curve Turbidity – Biomass (as VSS) for microbiome R1.

**Table A1.** Acetate consumption (in %) of each microbiome across the different trials performed. Note that in trial 9, no Ac was added to the medium (see Table 2 from the main text).

| | Microbiome | | | | | | |
|---|---|---|---|---|---|---|---|
| **Trial** | **UP** | **R1** | **R2** | **R3** | **CC** | **CW1** | **CW2** |
| 1 | 44.8 | 48.0 | 43.9 | 66.3 | 72.8 | 63.7 | 36.5 |
| 2 | 67.1 | 62.9 | 60.5 | 63.9 | 66.5 | 54.3 | 41.0 |
| 3 | 55.2 | 57.8 | 53.5 | 50.0 | 100 | 72.7 | 42.3 |
| 4 | 66.6 | 36.0 | 34.7 | 46.0 | 100 | 99.3 | 55.0 |
| 5* | 59.2 | 49.0 | 56.0 | 45.5 | 75.4 | 74.1 | 60.0 |
| 6 | 42.8 | 46.0 | 64.5 | 52.1 | 65.9 | 70.2 | 64.0 |
| 7 | 54.1 | 66.5 | 65.3 | 54.0 | 84.0 | 87.0 | 65.0 |
| 8 | 58.2 | 71.4 | 56.5 | 59.4 | 75.1 | 70.0 | 62.0 |
| 9 | - | - | - | - | - | - | - |

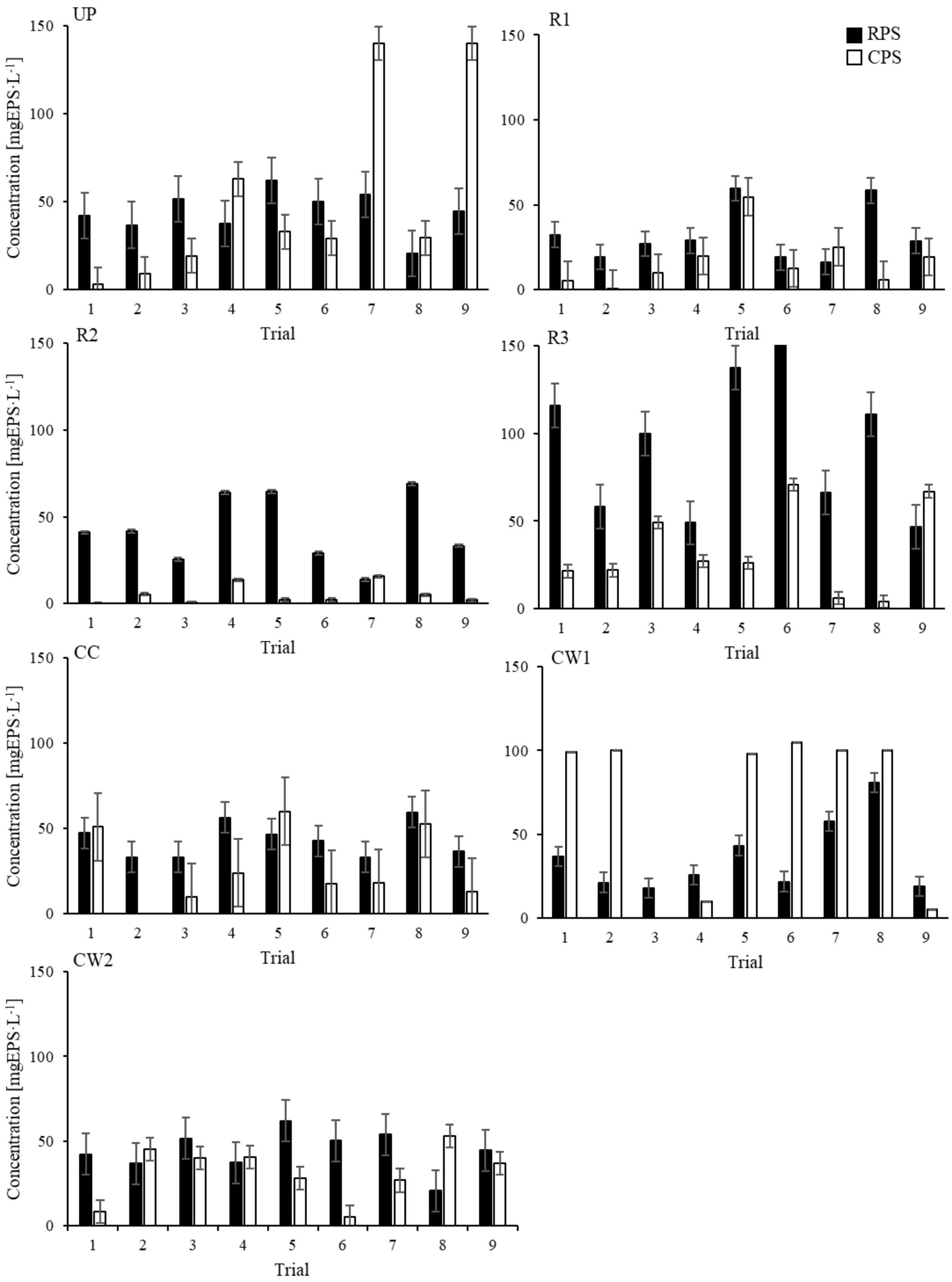


**Figure A4.** EPS production (RPS and CPS) for each microbiome under the nine tested trials (see **¡Error! No se encuentra el origen de la referencia.** for the properties of each trial). Standard deviation below 10 is not shown in the figure.

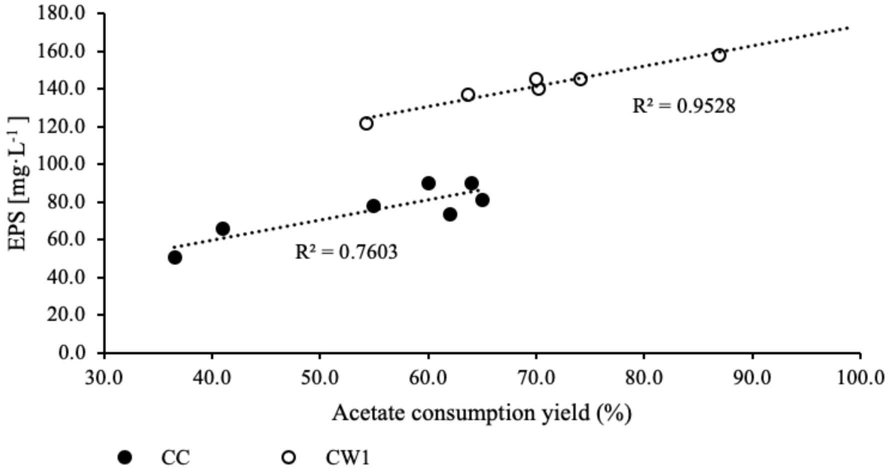


**Figure A5.** Correlation between acetate consumption (in %) and EPS concentration in microbiomes CW1 and CC.

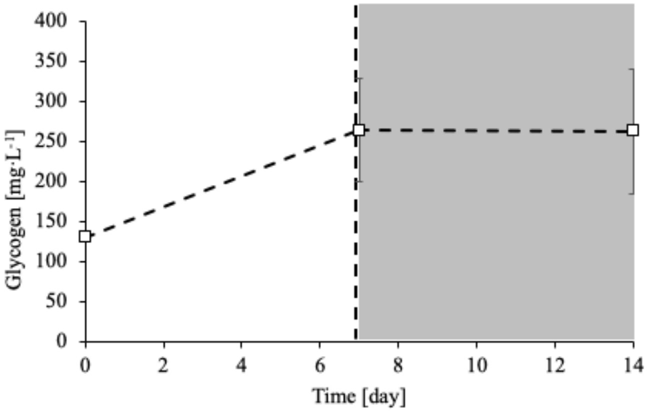


**Figure A6.** Concentration dynamics in Glycogen through the simultaneous PHB and EPS synthesis experiment.

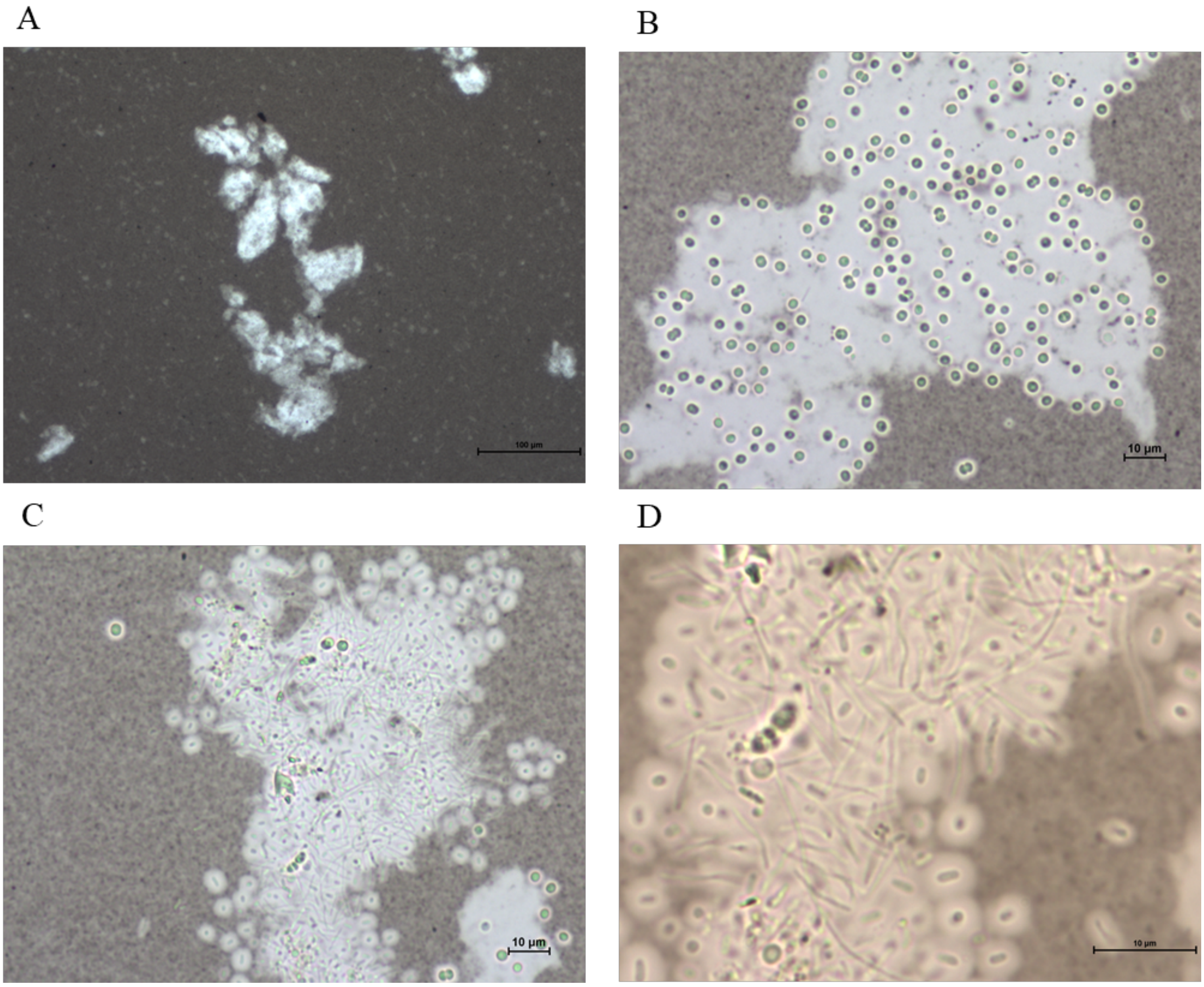


**Figure A7.** Light micrographs of microbiome R1 after black Chinese ink staining revealing EPS. (A) Image at 10X. (B) and (C) Show an aggregation of *Synechocystis* sp. and *Synechoccous* sp. in 40x. Filamentous bacteria were also present in some of the aggregates. (D) Detailed depiction of an aggregate at 100X.
